# Antibiotic resistance among *Escherichia coli* and *Salmonella* isolated from dairy cattle feces in Texas

**DOI:** 10.1101/2020.11.03.366336

**Authors:** Rosine Manishimwe, Paola M. Moncada, Marie Bugarel, H. Morgan Scott, Guy H. Loneragan

## Abstract

This study was conducted to develop and field-test a low cost protocol to estimate the isolate- and sample-level prevalence of resistance to critically important antibiotic drugs among *Escherichia coli* and *Salmonella* isolated from dairy cattle feces. *E. coli* and *Salmonella* were isolated from and screened on selective media, with and without antibiotics respectively. Bacterial isolates were further tested for susceptibility to a suite of antibiotics using disk diffusion. Molecular methods were performed on select bacterial isolates to identify and distinguish genetic determinants associated with the observed phenotypes. Among 85 non-type-specific *E. coli* randomly isolated from MacConkey agar without antibiotics, the isolate-level prevalence of resistance to tetracycline was the highest (8.2%), there was no isolate resistant to third-generation cephalosporin (0.0%) and one isolate was resistant to nalidixic acid (1.2%). Among 37 *E. coli* recovered from MacConkey agar with cefotaxime at 1.0µg/ml, 100% were resistant to ampicillin and 56.8% were resistant to a third-generation cephalosporin (ceftriaxone). Among 22 *E. coli* isolates recovered from MacConkey agar with ciprofloxacin at 0.5µg/ml, 90.9% were resistant to tetracycline whereas 77.3% and 54.5% were resistant to nalidixic acid and ciprofloxacin respectively. Sixteen *Salmonella* were isolated and only one demonstrated any resistance (i.e., single resistance to streptomycin). Among *E. coli* isolates that were either resistant or intermediate to ceftriaxone, an AmpC phenotype was more common than an extended spectrum beta-lactamase (ESBL) phenotype (29 versus 10 isolates, respectively). Among 24 *E. coli* isolates that were whole genome sequenced, phenotypic profiles of antibiotic resistance detected were generally substantiated by genotypic profiles. For instance, all isolates with an AmpC phenotype carried a *bla*_CMY2_ gene. The protocol used in this study is suited to detecting and estimating prevalence of antimicrobial resistance in bacteria isolated from food animal feces in resource-limited laboratories in the developing world.

## Introduction

Monitoring of the emergence, spread, and changes in levels of antimicrobial resistant bacteria along the food chain is needed to inform and guide integrated strategies for combating antimicrobial resistance[1]. In most cases, surveillance systems of antibiotic resistant bacteria in food animals target pathogenic bacteria, such as *Salmonella* and *Campylobacter* as well as indicator bacteria, such as *E. coli* and *Enterococcus spp*. After their isolation from samples, genus/species confirmation, and subtyping when necessary, the bacteria of interest are tested for susceptibility to a select number of antibiotics using a standard phenotyping method of choice. Even though the broth microdilution method is preferred in many surveillance systems and in research [2–4],other less technology intensive methods of antibiotic susceptibility testing, such as disk diffusion, may also be used[1,5]. After phenotypic antibiotic susceptibility testing, genetic characterization can be done through the detection of targeted genes or through whole genome sequencing (WGS). Additionally, the selective culture and detection of bacteria with rare antimicrobial resistance (AMR) mechanisms is highly recommended[1].

In many developed countries, AMR surveillance systems are replacing phenotypic antimicrobial susceptibility testing with WGS[6,7]. In many developing countries, however, surveillance systems are still not well established and the startup costs associated with equipment acquisition and maintenance can be prohibitive [8]. One of the main reasons impeding the establishment of AMR surveillance systems in these countries is the lack of sufficient resources needed for establishment, and then sustainment of the surveillance systems[8,9]. An inexpensive and reliable protocol that can generate sufficient high quality and reproducible information of the burden of antibiotic resistance among isolated bacteria would be one of the most helpful solutions for establishment of an AMR surveillance system in situation where resources are limited. Studies have demonstrated that the disk diffusion method is a cost effective method that can generate results comparable to other phenotypic methods of antibiotic susceptibility testing, that is, such as the broth microdilution or the agar dilution methods[10]. This method can be used efficiently to establish phenotypic antimicrobial resistance profiles and to detect mechanisms of resistance such as the production of extended spectrum beta-lactamases (ESBLs) among bacterial isolates.

We designed a protocol that uses the disk diffusion method to determine the isolate-level prevalence of resistant to various antibiotics and the sample-level prevalence of any bacteria not susceptible to third-generation cephalosporin or quinolone antibiotics. Additionally, the protocol described herein was designed to estimate the proportion of bacteria resistant to third-generation cephalosporins that produce either ESBL or AmpC enzymes. Afterward, the developed protocol was field-tested on fecal samples collected from dairy cattle to estimate isolate-level and sample-level prevalence of AMR among *E. coli* and isolate-level and sample-level prevalence among *Salmonella*.

## Materials and Methods

### Sample collection

Using a convenience sampling scheme, we collected 85 freshly voided fecal samples from dairy cattle of different age groups at a dairy farm located near Lubbock, Texas. Fecal samples were aseptically collected into sterilized polypropylene specimen containers then kept on wet ice and transported to a microbiology laboratory at Texas Tech University.

As fecal samples were collected from the pen-floor, there was no interaction with vertebrate animals, consequently an approval from an Institutional Animal Care and Use Committee wasn’t needed.

### Isolation of bacteria from fecal samples

In the laboratory, 10g of each fecal sample was weighted in a 710mL Whirl Pak® bag (Whirl-Pak, Madison, Wisconsin) and 90mL of buffered peptone water (Becton Dickinson, New Jersey, United States) was added. The mixture was placed in a commercial stomacher for 2 minutes at 230 rpm. Thereafter, the mixture was incubated at 42°C overnight prior to isolation of *E. coli* and *Salmonella*.

#### Isolation and identification of *Escherichia coli*

From each overnight non-selective enrichment, a 10μL loopful was streaked onto MacConkey agar (MAC, Hardy Diagnostics, California, United States) to isolate non-type-specific *E. coli*(NTS *E. coli*), meanwhile, another 10 μL loopful was streaked onto MAC supplemented with 1μg/mL of cefotaxime (MAC+CTX) to screen for *E. coli* resistant to third-generation cephalosporins (3GCr *E. coli*). An additional 10 μL loopful was streaked onto MAC supplemented with 0.5μg/mL of ciprofloxacin (MAC+CIP) to screen for *E. coli* not susceptible to quinolones (Qr *E. coli*). All three MacConkey agar plate types were incubated at 37°C overnight. Following the incubation, agar plates were inspected to identify growth of colonies with typical morphology of *E. coli* (i.e., pink, convex, circular and dry colonies with a surrounding pink zone). From each type of MacConkey agar plate, one typical colony was selected and re-streaked onto a similar MacConkey agar plate type for isolation of pure colonies. All well isolated presumptive *E. coli* were tested for indole production using an indole spot test (Hardy Diagnostics, California, United States) and were confirmed as *E. coli* by detection of the *wecA* gene using a real time polymerase chain reaction (rtPCR).

#### Isolation and identification of *Salmonella*

One mL of each overnight non-selective enrichment was transferred into 9mL of Rappaport-Vassiliadis *Salmonella* broth (Hardy Diagnostics, California, United States) and another 1mL was transferred into 9mL of tetrathionate broth (Hardy Diagnostics, California, United States). Both inoculated broths were incubated at 42°C overnight. After incubation, a 10μL loopful of each broth was streaked onto brilliant green sulfa agar (BGS, Becton Dickson, New Jersey, United States) and onto xylose lysine deoxycholate agar (XLD, Hardy Diagnostics, California, United States) to isolate *Salmonella*. In addition, 10 μL loopful of each broth was streaked onto BGS and onto XLD agar plates, each supplemented with 1μg/mL of cefotaxime (BGS+CTX and XLD+CTX) to screen for *Salmonella* resistant to third-generation cephalosporins. Another 10μL loopful of each broth was streaked onto BGS and XLD agar plates both supplemented with 0.5μg/mL of ciprofloxacin (BGS+CIP and XLD+CIP) to screen for *Salmonella* not susceptible to quinolones. All agar plates were incubated at 37°C overnight. Following incubation, agar plates were inspected for growth of colonies with morphology typical of *Salmonella* (i.e, pink, circular, dry, convex colonies on BGS; black, circular convex colonies on XLD). From each type of agar plate, a single typical colony was selected to be re-streaked onto the same type of agar plate for isolation of pure colonies. All presumptive *Salmonella* were tested for production of H_2_S gas, dextrose fermentation and decarboxylation reaction using lysine iron agar (Hardy Diagnostics, California, United States) and were confirmed to be *Salmonella* by detection of the *ttrC* gene using rtPCR.

### Antibiotic susceptibility testing

All isolates confirmed as *E. coli* and *Salmonella* were tested for susceptibility to a panel of 12 antibiotics using the disk diffusion method in accordance to guidelines of the Clinical Laboratory Standard Institute (CLSI)[11]. The antibiotics and concentration in each disk were amoxicillin-clavulanic acid 20/10μg(AMC), ampicillin 10μg (AMP), azithromycin 15μg (AZI), cefoxitin 30μg (FOX), ceftriaxone 30μg (CRO), chloramphenicol 30μg (CHL), ciprofloxacin 5μg (CIP), colistin 10μg (COL), meropenem 10µg (MER), nalidixic acid 30μg (NAL),streptomycin 10µg (STR), and tetracycline 30μg (TET). Inhibition zone diameters around the antibiotic-impregnated disks were measured in mm and rounded to the closest integer before being compared to the CLSI clinical break points in order to classify each bacterial isolate as resistant, intermediate or susceptible[11]. Because there was no CLSI standard inhibition zone diameters for colistin, these data were interpreted in accordance with a study conducted by Galani and collaborators[12].

Based on previous investigation of antimicrobial resistance in the region, all bacterial isolates not susceptible (i.e., intermediate and resistant) to a third-generation cephalosporin (ceftriaxone) were expected to have a phenotype reflecting ESBL- or AmpC-production. Suspected ESBL- or AmpC beta-lactamase-producing bacteria were discriminated by the combination disk test (CDT) according to CLSI guidelines[11] using a second panel of 12 antibiotic-impregnated disks. In addition to cefotaxime 30μg (CTX), cefotaxime-clavulanic acid 30/10μg (CTX-CLA), ceftazidime 30μg (CAZ), and ceftazidime-clavulanic acid 30/10μg (CAZ-CLA), the 4 antibiotic disks required by the CDT method, the second panel of antibiotics included amikacin 30μg (AMK),cefazolin 30μg (CFZ), cefepime 30μg, (FEP), fosfomycin 200μg (FOS), gentamicin 10μg (GEN), imipenem 10μg (IMP), sulfisoxazole 300μg (SSS), and trimethoprim/sulfamethoxazole 1.25/23.75μg (SXT).A bacterial isolate was classified to have a phenotype of ESBL-production when the absolute difference between the inhibition zone diameter around ceftazidime (coded as resistant) versus ceftazidime-clavulanic acid (CLA) and/or around cefotaxime (coded as resistant) versus cefotaxime-clavulanic acid was equal to or greater than 5mm. An isolate was classified to have a phenotype of AmpC beta-lactamase production when the difference between the inhibition zone diameter around ceftazidime (coded as resistant) versus ceftazidime-clavulanic acid and/or around cefotaxime (coded as resistant) versus cefotaxime-clavulanic acid was less than 5mm[11].

*E*.*coli* ATCC 25922 and *Klebsiella pneumoniae* ATCC 700603 were used as quality control strains.

### Polymerase chain reaction to detect genes encoding for beta-lactamases

All isolates confirmed as exhibiting either a phenotype suggestive of ESBL or else AmpC beta-lactamase-production were subjected to DNA extraction using a boiling preparation method. The extracted DNA was used as a template to detected the family of *bla_CTX-M_* genes (genes encoding for ESBL production) or *bla*_CMY-2_ gene (gene encoding for AmpC beta-lactamase production) by conventional PCR (cPCR), using previously published primers[13].

### Whole genome sequencing (WGS)

Following phenotypic AMR characterization, 24 *E. coli* isolates classified as either ESBL or AmpC phenotypes, resistant to NAL or reduced susceptibility to CIP were selected for WGS. The DNA was extracted using a commercial DNA extraction kit (Qiagen,Venlo, Netherland), libraries were prepared using the Nextera XT DBA library preparation kit (Illumina, California, United States) and the sequencing was performed using an Illumina Miseq (Illumina, California, United States). Generated raw reads (fastq files) were assembled using SPAdes 3.9 on the Center for Genomic Epidemiology platform.

### Data analysis

The prevalence proportion (expressed as percentage) of bacteria resistant to each tested antibiotic was determined and confidence intervals were calculated as 95% binomial proportions representing Wilson intervals using R.3.0. software. The proportion (expressed as percentages) of samples with bacteria not susceptible (i.e., resistant or intermediate) to a third-generation cephalosporin (ceftriaxone) or else to quinolones (nalidixic acid and/or ciprofloxacin) was calculated by dividing the number of samples with non-susceptible bacteria by the total number of collected samples collected (i.e., only those isolates screened on media with antibiotics were used to calculate the sample-level percentages). Whole genome sequencing data were analyzed using various bioinformatic tools found on the website of the Center for Genomic Epidemiology, including Resfinder3.0 that detect mobilizable genes and chromosomal mutations conferring antibiotic resistance in bacteria.

## Results

### Isolated bacteria

In total, NTS *E. coli* were recovered from all 85 fecal samples. The recovery rate of *Salmonella* and of either bacterial species (i.e., *E. coli* or *Salmonella enterica*) presumptively resistant to third-generation cephalosporins or else resistant/reduced susceptibility to quinolones was lower (Table1).

**Table 1.** Numbers of bacteria isolated from dairy cattle feces at a dairy farm in Texas.

| Bacteria | Culture medium of isolation | #of samples | # of isolates | % |
| --- | --- | --- | --- | --- |
| ***E. coli*** |  |  |  |  |
| NTS *E. coli* | MAC | 85 | 85 | 100.0% |
| Pres. 3GCr *E. coli* | MAC + CTX | 85 | 37 | 43.5% |
| Pres. Qr *E. coli* | MAC + CIP | 85 | 22 | 25.9% |
| ***Salmonella*** |  |  |  |  |
| *Salmonella* | BGS or XLD | 85 | 16 | 18.8% |
| Pres. 3GCr *Salmonella* | BGS + CTX or XLD + CTX | 85 | 1 | 1.2% |
| Pres. Qr *Salmonella* | BGS + CIP or XLD + CIP | 85 | 0 | 0.0% |

#: number; MAC: MacConkey agar; BGS: brilliant green sulfa; XLD: xylose lysine deoxycholate. MAC+CTX, BGS+CTX, XLD+CTX: respective culture medium supplemented with 1 µg/mL of cefotaxime. MAC+CIP, BGS+CIP, XLD+CIP: respective culture medium supplemented with 0.5 µg/mL of ciprofloxacin. Pres.3GCr: presumptive third-generation cephalosporin resistant, Pres. Qr: Presumptive quinolone resistant (or reduced susceptibility).

### Antibiotic resistance among isolated bacteria

Antibiotic susceptibility testing of isolated bacteria showed that resistance to antibiotics was rare among NTS *E. coli* isolated on MAC when compared to presumptive 3GCr *E. coli* screened on MAC+CTX or else presumptive Qr *E. coli* screened on MAC+CIP. Isolate-level prevalence of resistance to tetracycline (8.2%) was the highest among NTS *E. coli* isolates while resistance to cefoxitin, colistin, meropenem, ceftriaxone, and ciprofloxacin were completely absent among these bacterial isolates (Table 2).

**Table 2.**
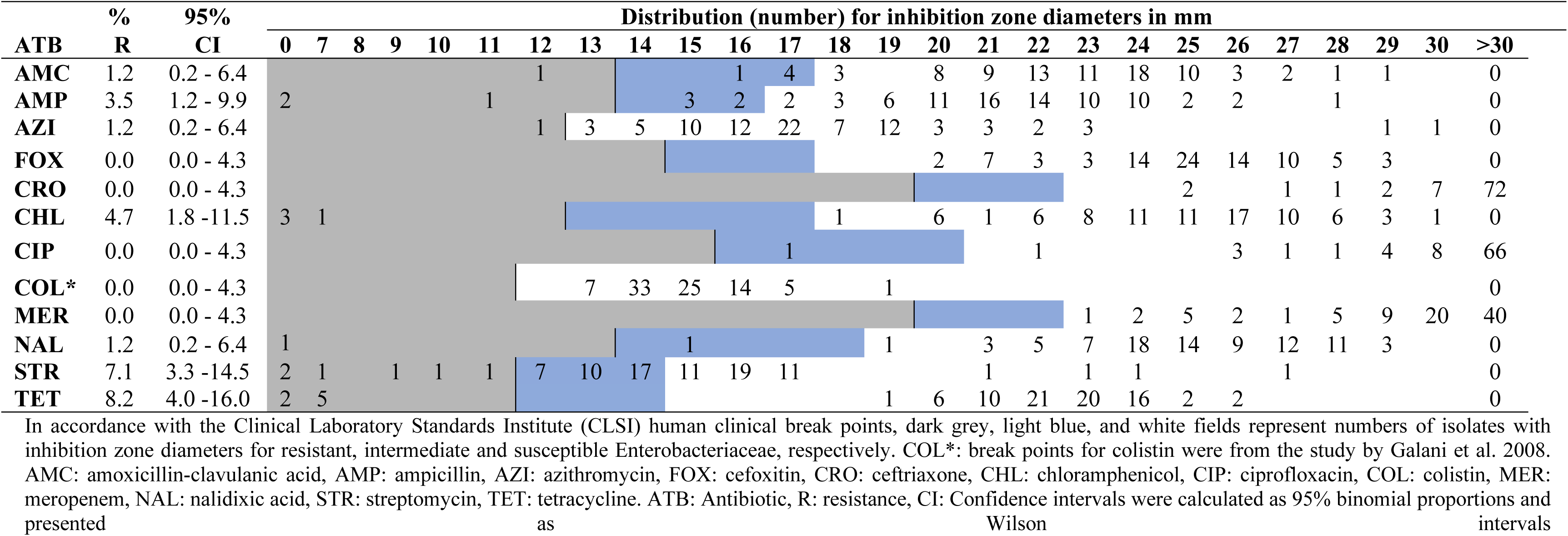
Distribution of inhibition zone diameters of non-type-specific *E. coli* (n=85) isolated on plain MacConkey agar (without antibiotic)

Among the 37 presumptive 3GCr *E. coli* screened on MAC+CTX, all isolates were resistant to ampicillin (100%), 21 isolates were resistant to ceftriaxone (55.8%), and none of the isolates was resistant to meropenem (Table 3). In total, 36 out of the 37 presumptive 3GCr *E. coli* were not susceptible (i.e., either resistant or intermediate) to a third-generation cephalosporin (CRO). These isolates were from 36 out of the 85 collected samples. The sample-level prevalence of *E. coli* non-susceptible to third-generation cephalosporin was calculated to be 42.3% (36/85) with a 95% confidence interval of 32.4% - 53.0%.

**Table 3.**
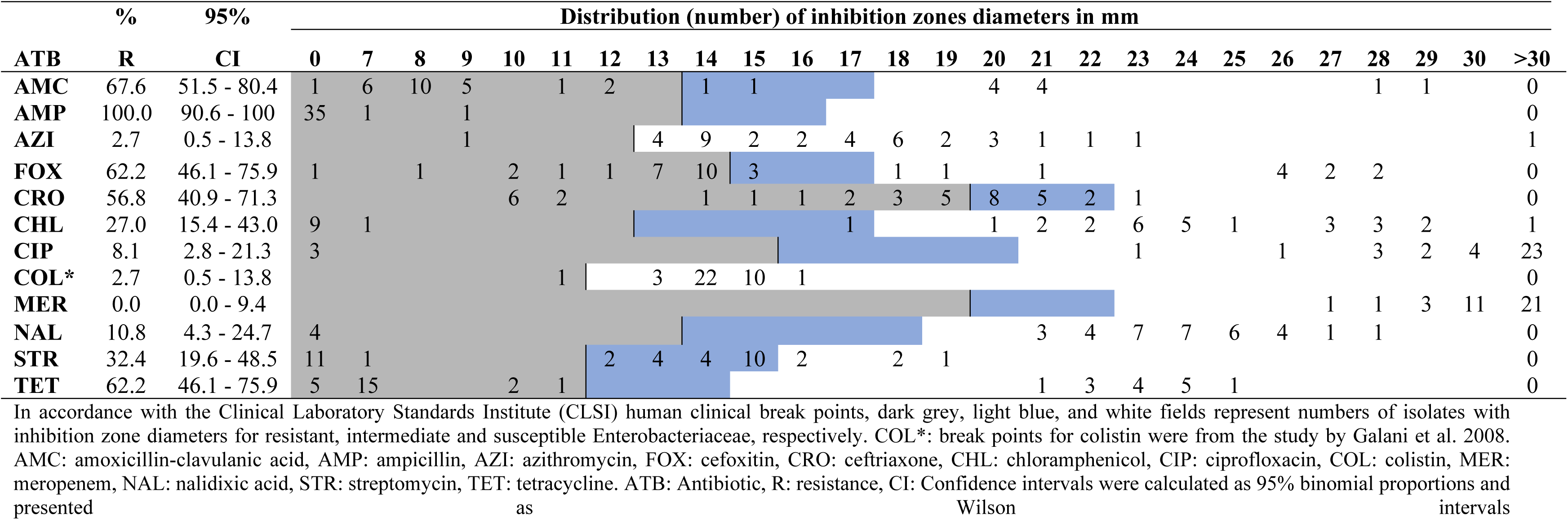
Distribution of inhibition zone diameters of presumptive third-generation cephalosporin resistant *E. coli* (n=37) isolated on MacConkey agar supplemented with 1 µg/mL cefotaxime.

Most presumptive Qr *E. coli* screened on MAC+CIP were resistant to tetracycline (90.9%), meanwhile 17 isolates were resistant to nalidixic acid (77.3%), 12 isolates were resistant to ciprofloxacin (54.5%), and none of the isolates was resistant to colistin or meropenem (Table4). All of the 22 presumptive Qr *E. coli* were resistant or intermediate to nalidixic acid or else to ciprofloxacin. These isolates were recovered from 22 out of 85 collected samples. The sample-level prevalence of *E. coli* non-susceptible to quinolone antibiotics was calculated to be 25.9 % with a 95% confidence intervals of 17.8%-36.1%. In total 12 out of the 22 Qr *E. coli* isolates were resistant to both nalidixic acid and ciprofloxacin, 4 isolates were resistant to nalidixic acid only, 5 isolates were intermediate to nalidixic acid but susceptible to ciprofloxacin and 1 isolate was intermediate to both ciprofloxacin and nalidixic acid.

**Table 4.**
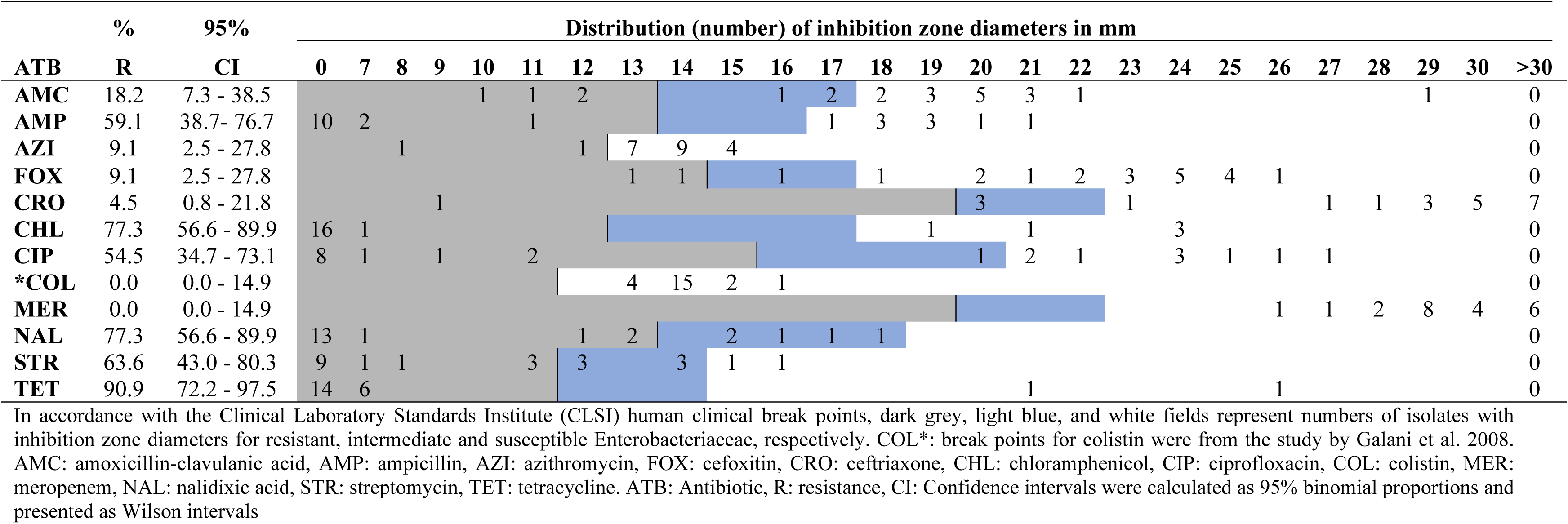
Distribution of inhibition zone diameters of presumptive quinolone resistant *E. coli* (n=22) isolated on MacConkey agar supplemented with 0.5 µg/mL of ciprofloxacin

Resistance to antibiotics among *Salmonella* isolated on culture media without antibiotics was very low yielding a single isolate that was resistant to only streptomycin. The sole *Salmonella* isolated on XLD+CTX was confirmed to be resistant to CRO.

### Phenotypic and genotypic detection of ESBL- and AmpC-producing bacteria

The sole *Salmonella* isolated from XLD+CTX plates, exhibited the phenotype of an AmpC beta-lactamase producer; later, this was confirmed by the presence of *bla*_CMY-2_gene using cPCR. A total of 40 *E. coli* isolates were found to be resistant or intermediate to ceftriaxone.

Thirty-six of them were isolated from MAC+CTX and four were isolated from MAC+CIP plates. The second panel of antibiotics showed that these isolates were resistant to at least one of the third-generation cephalosporins tested; furthermore all of them were resistant to cefazolin. Only a very few of these *E. coli* isolates were resistant to sulfisoxazole and trimethoprim/sulfamethoxazole (Table 5).

**Table 5.**
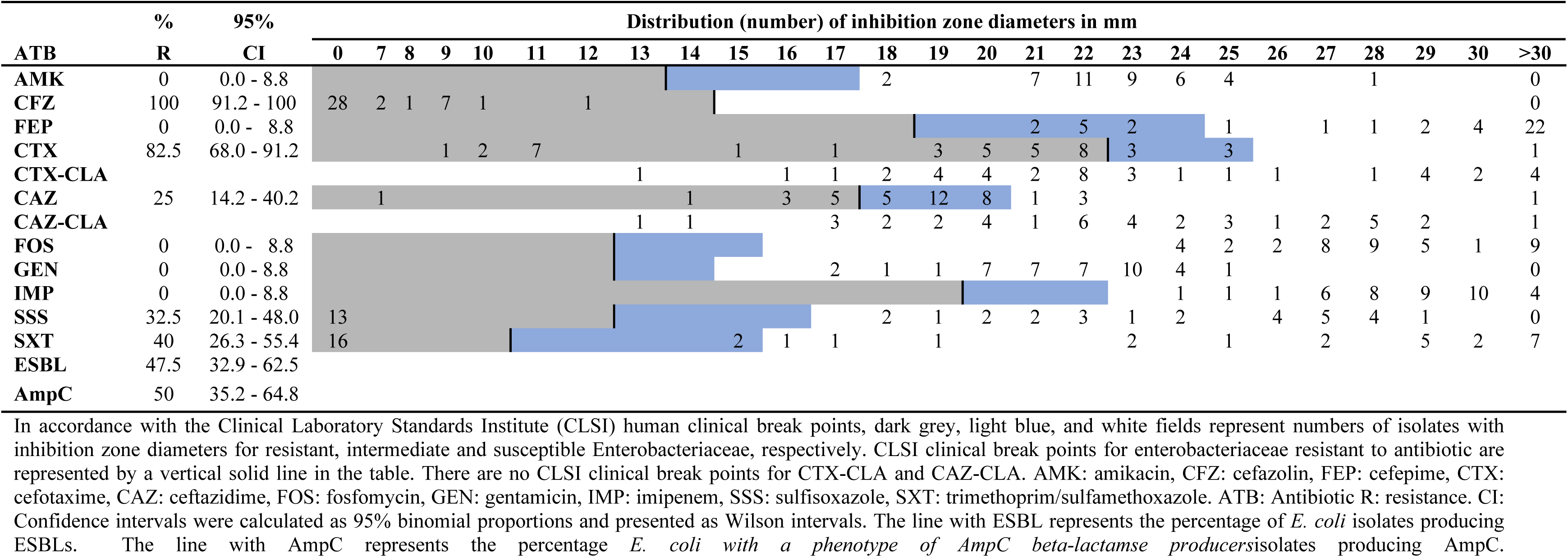
Distribution of inhibition zone diameters of all *E. coli* isolates (n=40) not susceptible to third-generation cephalosporins (i.e., second panel of antibiotics)

The phenotypic combination disk test detected 19 *E. coli* isolates with a phenotype indicating ESBL-production, 20 isolates with a phenotype indicating AmpC beta-lactamase-production and a single isolate that was not confirmed to be resistant to either cefotaxime or ceftazidime. On the other hand, molecular cPCR detected genes encoding for ESBL production (*bla*_*CTX-M-1*_or *bla*_*CTX-M-9*_ family genes) in 10 isolates (25.0%), the gene encoding for AmpC beta-lactamase production (i.e., *bla*_CMY-2_) in 29 isolates (72.5%) and none of these genes in one isolate (2.5%).

### Whole genome sequencing of select *E. coli* isolates

Whole genome sequencing of 24 *E. coli* isolates showed that the gene *tet*(A) encoding for a tetracycline efflux pump, was present in 70.8% of sequenced isolates. In general, all the detected genes were in accordance with the phenotypic antibiotic resistance observed in each of the isolate tested. Some exception included the presence of the *aph (3’)-Ia* gene that purportedly confers resistance to aminoglycosides, though in our case, in isolates phenotypically susceptible to these antibiotics. The mutation *gyrA[87:D-Y]* that confers resistance to quinolone antibiotics was observed in one isolate exhibiting no phenotypic resistance to either nalidixic acid or ciprofloxacin. The presence of the *mef(B)* gene that encodes for resistance to macrolides was detected in one isolate susceptible to azithromycin. Finally, the mutation *pmrB[161: V-G]* that confers resistance to polymixins was observed in an isolate phenotypically susceptible to colistin (Fig. 1).

**Fig. 1.**
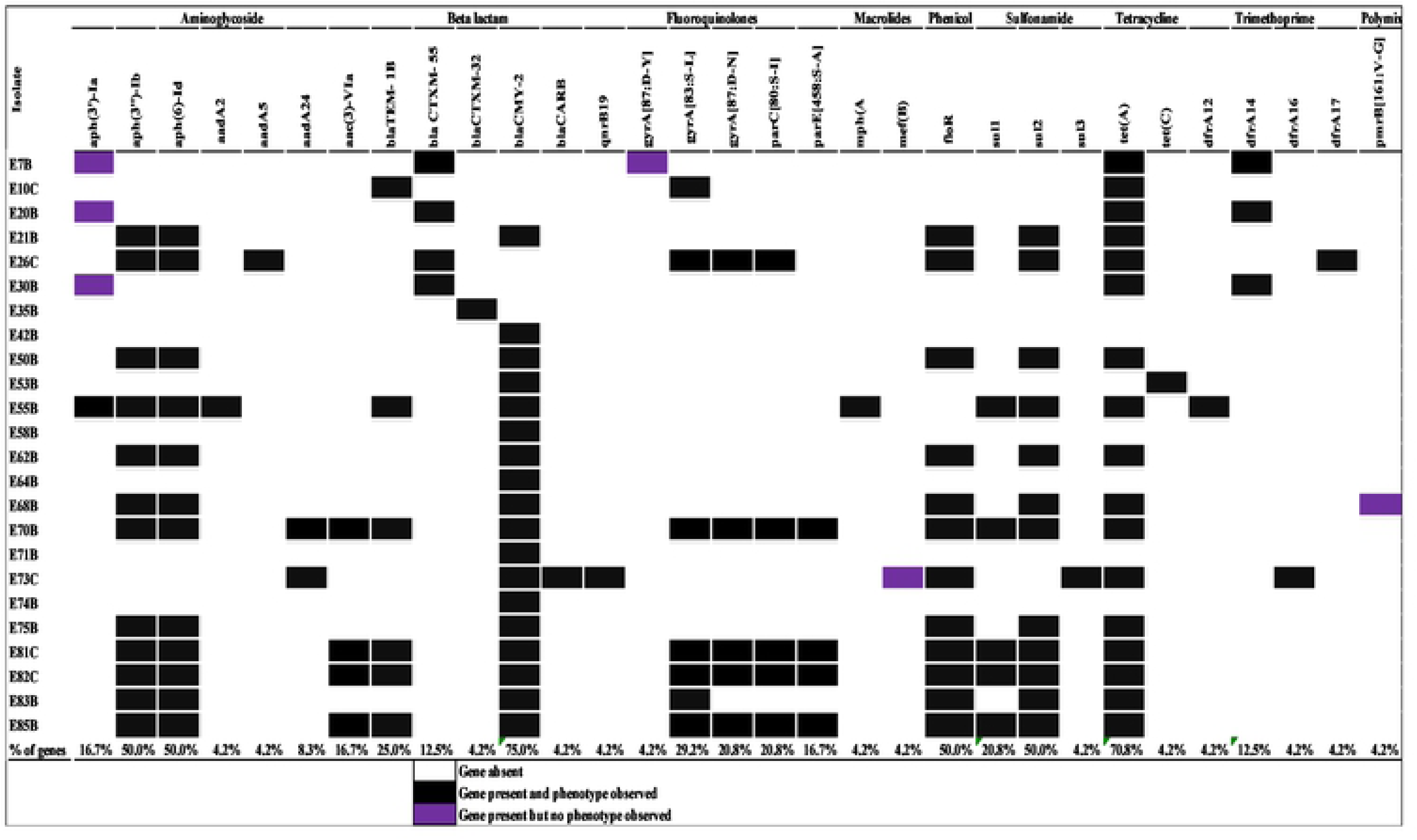
Genetic determinants of antibiotic resistance among *E. coli* isolated from cattle feces on a dairy farm in Texas. Black cells indicate the presence of a resistance determinant and a phenotype of that determinant in a bacterial isolate. Purple cells indicate the presence of a resistance determinant without a corresponding phenotype in the same isolate. White cells indicate the absence of both resistance determinant and corresponding phenotype.

Finally, WGS revealed that *E. coli* isolates resistant to nalidixic acid typically had only a single point mutation in the *gyrA* gene or else harbored a plasmid-mediated quinolone-resistance gene (*qnr*). All isolates resistant to both nalidixic acid and ciprofloxacin had a mutation in both *gyrA* and *parE* or *parC* genes of the quinolone resistance determining region (QRDR).

## Discussion

This study was conducted to field-test a cost-effective and highly valid protocol that could be used to determine the status of antibiotic resistance among *E. coli* and *Salmonella* isolated from food-producing animals, especially where laboratory resources are limited. The protocol used in this study was inspired by different guidelines for antimicrobial resistance detection, including guidelines from the European Food Safety Authority(EFSA)[14], the U.S National Antimicrobial Resistance Monitoring System (NARMS)[2], the European Union Reference Laboratory for Antimicrobial Resistance (EURL-AR)[15], and the Danish Integrated Antimicrobial Resistance Monitoring and Research Program (DANMAP)[16]. Unlike the above mentioned guidelines, this protocol used the disk diffusion method instead of broth or agar dilution methods because disk diffusion is recognized as a simple and low-cost method when compared to other antibiotic susceptibility testing methods[17]. To increase our confidence in recommending the protocol, results of the phenotypic methods used herein were thereafter cross-referenced and validated using by results from relevant molecular methods.

In regard to results obtained in this study, all NTS *E. coli* and *Salmonella* isolated on bacterial culture media without supplemented antibiotics were largely susceptible to all antibiotics tested; meanwhile, bacteria isolated on culture media with antibiotics supplemented at sub-breakpoint levels tended to be resistant to more than three antibiotics. This observation provided evidence that when a bacterium acquires resistance to one antibiotic it tends also to be resistant to other antibiotics. In fact, in most of the cases, different genes encoding antibiotic resistance are known to be co-located on transmissible genetic elements such as plasmids. When a resistance plasmid is transferred to a previously susceptible bacterium, multidrug resistance can be transferred in a single conjugation event[18].

Furthermore, percentages of samples that generated presumptive 3GCr *E. coli* (43.5%) and presumptive 3GCr *Salmonella* (1.2%) were lower than the percentages of samples with presumptive 3GCr *E. coli* (89.1%) and presumptive 3GCr *Salmonella* (10.9%) reported in three beef feed lots in Nebraska[19]. In addition, similar to the study in Nebraska[19], we also noted that the number of 3GCr *E. coli* isolates phenotypically or genotypically confirmed to be AmpC-producers was higher than the number of isolates confirmed to be ESBLs-producers. The only identified 3GCr *Salmonella* was confirmed to be an AmpC beta-lactamase producer. In the U.S, resistance to third-generation cephalosporins among *Salmonella* from food animals has historically been largely due to the gene ^*bla*^CMY-2 encoding for AmpC beta-lactamase production [20].

In the present study, nine 3GCr *E. coli* had a phenotype typical of ESBL-production (according to the CDT) while the cPCR showed that these isolates carried *bla*_CMY-2_, a gene encoding for AmpC beta-lactamase production, instead of genes encoding for an ESBL. The combination disk test is reported to be an accurate phenotypic method and is widely used to detect ESBL-producing Enterobacteriaceae[21]. Despite its widespread use and solid reputation, the sensitivity and specificity of this method to detect ESBL-producing Enterobacteriaceae is not always 100%[22]. In fact, different authors have reported a number false positive ESBL-producing Enterobacteriaceae by the CDT in their investigations [23–26].

To mitigate misclassification of bacterial isolates as ESBL-producers using phenotypic methods, some of these authors have suggested modifications of standard methods, such as the CDT, in order to increase their efficacy in discriminating ESBL-producing bacteria from AmpC beta-lactamase-producing bacteria[23,24]. Along this line, our study illustrates that when the combination disk test is used to identify ESBs-producing *E. coli*, some precaution should be taken as the test may produce several false positive *E. coli* producing ESBLs, especially when the results are close to 5 mm decision point. A close look at the 9 *E. coli* isolates falsely classified as ESBL-producers showed that the difference between inhibition zone diameters around cefotaxime with clavulanic acid and around cefotaxime was less than 5mm for all the isolates. In contrast, the difference between inhibition zone diameters around ceftazidime with clavulanic acid and around ceftazidime was equal to 5mm or slightly higher than 5mm.The later observation led to the conclusion of classifying these same isolates as ESBL-producers. Importantly, 8 of the 9 isolates were not susceptible to cefoxitin (a second-generation cephalosporin (cephamycin) antibiotic).

Based on observations made in this study, we came up with the following rule of thumb: false positive ESBL-producing *E. coli* can be detected by looking at the increase in inhibition zone diameters around CAZ-CLA versus CAZ and around CTX-CLA versus CTX. False positive ESBL-producing *E. coli* can be identified as isolates for which inhibition zone diameters around ceftazidime is increased by 5mm or slightly higher (6mm) due to clavulanic acid while the increase of the inhibition zone diameter around cefotaxime caused by clavulanic acid is less than 5mm. When false positives are identified, we recommend using the information generated by CTX with and without CLA to classify an isolate as ESBL- or AmpC-producer. In addition, a look at bacterial isolates’ susceptibility to cefoxitin (a cephamycin) or amoxicillin-clavulanic acid maybe helpful to conclude that an isolate is not an ESBL-producer but might be an AmpC-producer. It is reported that AmpC-producing Enterobacteriaceae are resistant to cephamycins such as cefoxitin while ESBL-producers are susceptible to cephamycins[27,28].

In general, WGS showed that phenotypic antibiotic resistance observed in *E. coli* isolates was a good indicator of the presence of genetic antibiotic resistance determinants, with only few a exceptions. The observed discordance may be explained by the fact that the presence of a resistance gene doesn’t always mean its expression. In addition, some genes are cryptic or even misclassified as to their primary purpose in bacterial host function. Results of WGS also showed that all isolates resistant to both NAL and CIP had mutations in both genes of the quinolone resistance determining region (QRDR) while isolates resistant to NAL alone had a single mutation in one of the target genes, else or carried a plasmid mediated quinolone-resistance gene (*qnr* gene). In fact, mutations in the QRDR have been identified as the common genetic determinant conferring a higher level of resistance to quinolone antibiotics while PMQR genes confer moderate resistance[29,30]. Several studies have proven that the detection of genetic determinants of antibiotic resistance by WGS accurately predicts antibiotic resistance phenotypic behavior of bacterial isolates[6,31]. High level of agreement between WGS and phenotypic antibiotic susceptibility testing have been reported [32,33].

In conclusion, this study established that bacteria resistant to antibiotics were present in dairy cattle on the study farm but at a low level. The addition of an antibiotic to the culture medium of isolation helped in detecting and later characterizing the few antibiotic resistant bacteria. The developed protocol can help to establish percentages of indicator *E. coli* resistant to various antibiotics, including critically important antibiotics for human medicine, at a relatively low cost and with high reliability, even in the developing world. Furthermore, cPCR and WGS both supported our phenotypic findings at a high level which increased our confidence in recommending the tested protocol for its use to establish status of AMR in food animals; specifically, when laboratory facilities are limited and financial and other resources are scarce.

## Acknowledgements

We acknowledge the dairy farm management that allowed us to collect fecal samples.

## References

1. World Health Organization (WHO). Integrated Surveillance of Antimicrobial Resistance in Foodborne Bacteria: Application of a One Health Approach [Internet]. Geneva; 2017. doi:Licence: CC BY-NC-SA 3.0 IGO

2. The National Antimicrobial Resistance Monitoring System. The National Antimicrobial Resistance Monitoring System Manual of Laboratory Methods [Internet]. 2016. Available: https://www.fda.gov/downloads/AnimalVeterinary/SafetyHealth/AntimicrobialResistance/NationalAntimicrobialResistanceMonitoringSystem/UCM528831.pdf

3. Government of Canada. Canadian Integrated Program for Antimicrobial Resistance Surveillance (CIPARS) 2016 Annual Report [Internet]. Guelph, Ontario; 2018. Available: http://publications.gc.ca/collections/collection_2018/aspc-phac/HP2-4-2016-eng.pdf

4. National Veterinary Assay Laboratory. Report on the Japanese Veterinary Antimicrobial Resistance Monitoring System 2014-2015 [Internet]. Tokyo; 2018. Available: http://www.maff.go.jp/nval/yakuzai/yakuzai_p3.html

5. European Union Food Safety Authority. ECDC/EFSA/EMA second joint report on the integrated analysis of the consumption of antimicrobial agents and occurrence of antimicrobial resistance in bacteria from humans and food-producing animals. EFSA J. 2017;15. doi:10.2903/j.efsa.2017.4872

6. Neuert S, Nair S, Day MR, Doumith M, Ashton PM, Mellor KC, et al. Prediction of phenotypic antimicrobial resistance profiles from whole genome sequences of non-typhoidal Salmonella enterica. Front Microbiol. 2018;9: 1–11. doi:10.3389/fmicb.2018.00592

7. United States Department of Agriculture. The National Antimicrobial Resistance Monitoring System [Internet]. 2018 [cited 5 Feb 2019]. Available: https://pregunteleakaren.gov/wps/portal/fsis/topics/regulatory-compliance/!ut/p/a1/rVZLc5swGPwtOfioQUIP0NH1xG7tDq6dOI25ZARIVBleAZpp-usrnHbixA3CGcQBhHZX7Dc7n3BC59YJC_GoU9HqshBZNw_ZHdxAhvgMLiEy15cAX5HPywBDFxvA_hiw5mhuADeb9Wo2g36AX_PX82m3jOc3frBAkKBHfGFPby

8. Seale AC, Gordon NC, Islam J, Peacock SJ, Scott JAG. AMR Surveillance in low and middle-income settings -A roadmap for participation in the Global Antimicrobial Surveillance System (GLASS). Wellcome Open Res. 2017;2: 92. doi:10.12688/wellcomeopenres.12527.1

9. Ayukekbong JA, Ntemgwa M, Atabe AN. The threat of antimicrobial resistance in developing countries: Causes and control strategies. Antimicrob Resist Infect Control. Antimicrobial Resistance & Infection Control; 2017;6: 1–8. doi:10.1186/s13756-017-0208-x

10. Hoelzer K, Cummings KJ, Warnick LD, Schukken YH, Siler JD, Gröhn YT, et al. Agar Disk Diffusion and Automated Microbroth Dilution Produce Similar Antimicrobial Susceptibility Testing Results for Salmonella Serotypes Newport, Typhimurium, and 4,5,12:i-, But Differ in Economic Cost. Foodborne Pathog Dis. 2011;8: 1281–1288. doi:10.1089/fpd.2011.0933

11. Clinical and Laboratory Standards Institute. M100 Performance Standards for Antimicrobial Susceptibility Testing. 28th ed. Clinical and Laboratory Standards Institute. Pennsylvania, USA: Clinical and Laboratory Standards Institute; 2018.

12. Galani I, Kontopidou F, Souli M, Rekatsina P, Koratzanis E, Deliolanis J, et al. Colistin susceptibility testing by Etest and disk diffusion methods. Int J Antimicrob Agents. 2008;31:434–439. doi:10.1016/j.ijantimicag.2008.01.011

13. Dallenne C, Da Costa A, Decré D, Favier C, Arlet G. Development of a set of multiplex PCR assays for the detection of genes encoding important β-lactamases in Enterobacteriaceae. J Antimicrob Chemother. 2010;65: 490–495. doi:10.1093/jac/dkp498

14. European Food Safety Authority (EFSA), Aerts M, Battisti A, Hendriksen R, Kempf I C T, et al. Technical specifications on harmonised monitoring of antimicrobial resistance in zoonotic and indicator bacteria from food-producing animals and food. EFSA J. 2019;17: 122p. doi:10.2903/j.efsa.2019.5709

15. Henrik Hasman, Agersø Y, Hendriksen R, Cavaco LM, Guerra-Roman B. Isolation of ESBL-, AmpC-and carbapenemase-producing E. coli from caecal samples. Version 7. Eur Union Ref Lab Antimicrob Resist (EURL-AR). 2019; doi:10.2903/j.efsa.2011.2322.OJ

16. Danish Integrated Antimicrobial Resistance Monitoring and Research Programme (DANMAP). DANMAP 2018 - Use of antimicrobial agents and occurrence of antimicrobial resistance in bacteria from food animals, food and humans in Denmark [Internet]. Copenhagen; 2019. Available: www.danmap.org

17. Balouiri M, Sadiki M, Ibnsouda SK. Methods for in vitro evaluating antimicrobial activity: A review. J Pharm Anal. Elsevier; 2016;6: 71–79. doi:10.1016/j.jpha.2015.11.005

18. Hiroshi N. Multidrug Resistance in Bacteria. Annu Rev Biochem. 2010;78: 119–146. doi:10.1146/annurev.biochem.78.082907.145923.Multidrug

19. Schmidt JW, Agga GE, Bosilevac JM, Brichta-Harhay DM, Shackelford SD, Wang R, et al. Occurrence of antimicrobial-resistant Escherichia coli and Salmonella enterica in the beef cattle production and processing continuum. Appl Environ Microbiol. American Society for Microbiology; 2015;81: 713–725. doi:10.1128/AEM.03079-14

20. The National Antimicrobial Resistance Monitoring System. NARMS integrated report, 2015 [Internet]. Laurel, MD; 2017. Available: https://www.fda.gov/AnimalVeterinary/SafetyHealth/AntimicrobialResistance/NationalAntimicrobialRe

21. European Committee on Antimicrobial Susceptibility Testing. EUCAST guidelines for detection of resistance mechanisms and specific resistances of clinical and/or epidemiological importance. Versio 2.0/2017 [Internet]. The European Committee on Antimicrobial Susceptibility Testing. 2017. Available: http://www.eucast.org/fileadmin/src/media/PDFs/EUCAST_files/Resistance_mechanisms/EUCAST_detection_of_resistance_mechanisms_170711.pdf

22. Drieux L, Brossier F, Sougakoff W, Jarlier V. Phenotypic detection of extended-spectrum β-lactamase production in Enterobacteriaceae: Review and bench guide. Clin Microbiol Infect. European Society of Clinical Microbiology and Infectious Diseases; 2008;14: 90–103. doi:10.1111/j.1469-0691.2007.01846.x

23. Garrec H, Drieux-Rouzet L, Golmard JL, Jarlier V, Robert J. Comparison of nine phenotypic methods for detection of extended-spectrum β-lactamase production by enterobacteriaceae. J Clin Microbiol. 2011;49: 1048–1057. doi:10.1128/JCM.02130-10

24. Poulou A, Grivakou E, Vrioni G, Koumaki V, Pittaras T, Pournaras S, et al. Modified CLSI extended-spectrum β-lactamase (ESBL) confirmatory test for phenotypic detection of ESBLs among Enterobacteriaceae producing various β-lactamases. J Clin Microbiol. 2014;52: 1483–1489. doi:10.1128/JCM.03361-13

25. Polsfuss S, Bloemberg G V., Giger J, Meyer V, Böttger EC, Hombach M. Evaluation of a diagnostic flow chart for detection and confirmation of extended spectrum β-lactamases (ESBL) in Enterobacteriaceae. Clin Microbiol Infect. 2012;18: 1194–1204. doi:10.1111/j.1469-0691.2011.03737.x

26. Robberts FJL, Kohner PC, Patel R. Unreliable extended-spectrum β-lactamase detection in the presence of plasmid-mediated AmpC in Escherichia coli clinical isolates. J Clin Microbiol. 2009;47: 358–361. doi:10.1128/JCM.01687-08

27. Jacoby GA. AmpC B-Lactamases. Clin Microbiol Rev. 2009;22: 161–182. doi:10.1128/CMR.00036-08

28. Paterson DL, Bonomo RA. Clinical Update Extended-Spectrum Beta-Lactamases : a Clinical Update. Clin Microbiol Rev. 2005;18: 657–686. doi:10.1128/CMR.18.4.657

29. Correia S, Poeta P, Michel H. Mechanisms of quinolone action and resistance : where do we stand ? J Med Microbiol. 2017;66: 551–559. doi:10.1099/jmm.0.000475

30. Redgrave LS, Sutton SB, Webber MA, Piddock LJ V. Fluoroquinolone resistance: Mechanisms, impact on bacteria, and role in evolutionary success. Trends Microbiol. 2014;22: 438–445. doi:10.1016/j.tim.2014.04.007

31. Tyson GH, McDermott PF, Li C, Chen Y, Tadesse DA, Mukherjee S, et al. WGS accurately predicts antimicrobial resistance in Escherichia coli. J Antimicrob Chemother. 2015;70: 2763–2769. doi:10.1093/jac/dkv186

32. Shelburne SA, Kim J, Munita JM, Sahasrabhojane P, Shields RK, Press EG, et al. Whole-Genome Sequencing Accurately Identifies Resistance to Extended-Spectrum β-Lactams for Major Gram-Negative Bacterial Pathogens. Clin Infect Dis. 2017;65: 738–745. doi:10.1093/cid/cix417

33. Kumburu HH, Sonda T, Zwetselaar M Van, Leekitcharoenphon P, Lukjancenko O, Mmbaga BT, et al. Using WGS to identify antibiotic resistance genes and predict antimicrobial resistance phenotypes in MDR Acinetobacter baumannii in Tanzania 1–3. J Antimicrob Chemother. 2019;74: 1484–1493. doi:10.1093/jac/dkz055

